# Consumer risk perception towards pesticide-stained tomatoes in Uganda

**DOI:** 10.1101/2021.02.15.431249

**Authors:** Daniel Sekabojja, Aggrey Atuhaire, Victoria Nabankema, Deogratias Sekimpi, Erik Jors

## Abstract

**Background:** Tomatoes are consumed daily. Unfortunately, abuse of pesticide application by vegetable growers in Uganda increases risks of pesticide residue exposure among consumers, as they may be above Maximum Residue Limits (European Union Maximum Residue Limits used as a standard in Uganda). This study aimed to determine consumer attitudes and risk perceptions towards pesticide-stained tomatoes in Uganda to support interventions that could be used to reduce pesticide residue exposures in food.

**Methods:** A mixed methods cross-sectional study sampled 468 household consumers in four regions of Uganda, selecting one district (interventional project area) per region. In each district, about 60 household members were randomly selected from a total of three Sub Counties and interviewed. In addition, 9 tomato handlers (three tomato farmers, three tomato retailers, and three tomato wholesalers) participated in Focus Group Discussions (FGDs) per district. Collected data were entered into MS-Excel 13 and exported into Stata version 14.0 for cleaning and analysis at a 5% level of significance and 95% Confidence Intervals (CI). The proportion of risk perceptions and attitudes were computed and presented as percentages, while factors associated with risk perception were determined using Fisher exact test. Qualitative data collected under a traditional theory were analyzed using thematic content analysis.

**Results:** More than half, 54.2% (253/468), of the respondents were females, mean age was 37 years (SD=13.13, ranging from 18 to 88 years). Half of the respondents, 50.9% (238/467), were farmers by occupation, and 40.3% (188/468) had completed upper primary education. Only 5.0% (20/396) of consumers reported a high-risk perception towards tomatoes stained with pesticide residues, the rest, 95.0% (376/396), were buying pesticide-stained tomatoes despite their awareness of the possible health effects. The main reason for buying the pesticide-stained tomatoes was that a majority, 59.0% (230/390), lacked an alternative to stained tomatoes like organically grown tomatoes. However, consumers generally had a negative attitude towards pesticide-stained tomatoes, with 67.0% (313/468) of the consumers disagreeing with a statement that tomatoes sold on the market are safe. Consumer risk perception was significantly associated with their awareness about residues in the tomatoes; where the proportion of consumers who were aware of the risk of pesticide-stained tomatoes was 42.8 times more likely not to buy stained tomatoes compared to the proportion of those who were not aware. OR, 42.8 (95% CI: 10.76-170.28). However, after Fisher-Exact tests analysis, level of education P(0.975), gender P(0.581), and age group P(0.680) were not associated with consumer risk perception (95% CI and 5% level of significance).

**Conclusion:** Although the consumers had a negative attitude towards the pesticide-stained tomatoes, their risk perception towards them ranked low, with most consumers buying tomatoes stained with pesticide residues due to a lack of an alternative. Ministry of Agriculture extension service efforts should promote and emphasize community to start household-based organic kitchen gardens as the efforts for the establishment of a national pesticide residue monitoring center awaits.

## Introduction

Globally, there has been an increase in the inquiry on the knowledge of the dangers of chemicals in food which has aroused consumer concerns about food safety (1, 2). This follows from consumer reports on the health effects of pesticides from their inappropriate use, exposing consumers to high amounts of pesticide residues in harvested foods (3–5). Pesticide residues in food are directly related to the irrational application of pesticides on growing crops and a lesser extent, from residues remaining in the soil. Accumulated pesticide residues in food products that are absorbed in the human body are associated with human health hazards ranging from acute illnesses like; skin rashes, nausea, headaches, eye irritation, and shortness of breath to chronic toxic effects like; asthma, cancers, diabetes among other chronic illnesses (4, 6–8).

For numerous decades, pesticide use has intensified globally in agriculture, homes and industries aiming at increasing productivity and reducing losses (4, 9–12). In Sub-Saharan Africa, with a tropical climate that favors the growth and rapid multiplication of pests, pesticides are usually used at all levels of agricultural production, including on farms, to shield plants from pest attack and damage, to control weeds and parasites in livestock as well as in post-harvest control measures. It is now nearly impossible to produce food in tropical regions without using agrochemicals.

However, considerations have been made in the climate change mitigation strategies rolling on the adoption of agroecology mechanisms (13, 14). In developing countries, almost all fruits and vegetables grown commercially are sprayed with pesticides to combat pests and diseases. For example, a study done in 2014 at the two largest horticultural produce markets in Africa showed that 91% of the fruit and vegetable samples collected between 2012 and 2014 had pesticide residues, although these were compliant with the Maximum Residue Limits (MRLs) (15). A comparative study done in Uganda among two groups of farmers (organic vs conventional farmers) still attests to the fact that food consumers are still exposed to the pesticide through the consumption of contaminated foods and drinks apart from direct exposures during spraying (16).

In most low-income countries like Uganda, pesticide regulation enforcement and support of the agricultural extension staff to guide farmers on pesticide application standards and dosage is very minimal if not done by implementing partners like Non-Government Organizations. Unlike export products, fresh produce sold at local markets is not analyzed for agricultural chemical residues. This raises concerns about the perceived safety levels of local food supplies compared to exported products (15). For instance, a study in Uganda showed that 24.5% of farmers were not aware of any health risks of spraying tomatoes close to harvest time, almost 50% of farmers (45.8%) sprayed their tomatoes less than a week to harvest time, 29.2% sprayed their tomatoes on harvesting, with intentions to extend the shelf-life while 50% did so to attract consumers(17–19).

Another study in 2015, shows how farmers sprayed tomatoes 6 times the manufacturer recommended dosage and harvested these tomatoes 2-3 days after the last spraying session compared to the recommended pre-harvest interval of 4-7 days (18). These phytosanitary practices increase pesticide residues in tomatoes. This is further exacerbated by the lack of a pesticide residue monitoring plan for conventionally grown food and surveillance for pesticide poisoning in the health sector (20).

Although developed countries use 75% of global pesticides; these apply them with strict regulations compared to developing countries which lack regulation enforcement. Although developing countries use the least quantities of pesticides, they use the most toxic ones (21, 22) resulting in increased risks of acute poisoning. The inappropriate use of pesticides in developing countries increases pesticide exposure and health risks to consumers. Approximately 25% of developing countries lack regulations and 50% of the WHO-region countries lack sufficient resources to enforce their pesticide-related regulations (23, 24). Also, under existing international laws, highly toxic, banned or unregulated pesticides are always exported to developing countries (25–28), posing health risks to consumers.

Although Uganda is transitioning to establishing a pesticide residue monitoring program, this has moved slowly and made the protection of public health unrealized. There is limited published work in Uganda about consumer risk perception towards pesticides residues in food. This study, therefore, in its novelty, contributes to new knowledge in the areas of risk perception among Ugandan consumers through unveiling the understanding of consumers’ risk perception towards pesticide-stained tomatoes, and their attitudes towards pesticide-stained tomatoes to justify the need for a National Pesticide residue monitoring program in Uganda.

### Theoretical model

The study employed the risk perception model of consumer behavior, the most commonly used theoretical model for consumer risk perception. This model suggests that consumer risk perceptions are based on their cognitive, affective, and behavioral responses to potential risks associated with food. Cognitive responses include the consumer’s evaluation of the probability of a risk occurring and the severity of the potential impacts of the risk. Affective responses include the consumer’s emotional reactions to the risk, such as fear, anger, or disgust. Behavioral responses include the consumer’s decision to purchase or avoid the food in question, or to take other precautionary actions in response to the risk.

### Research questions

1. What is the risk perception of consumers regarding tomatoes stained with pesticides?
2. What are consumers’ attitudes towards tomatoes stained with pesticides?

### Conceptual framework

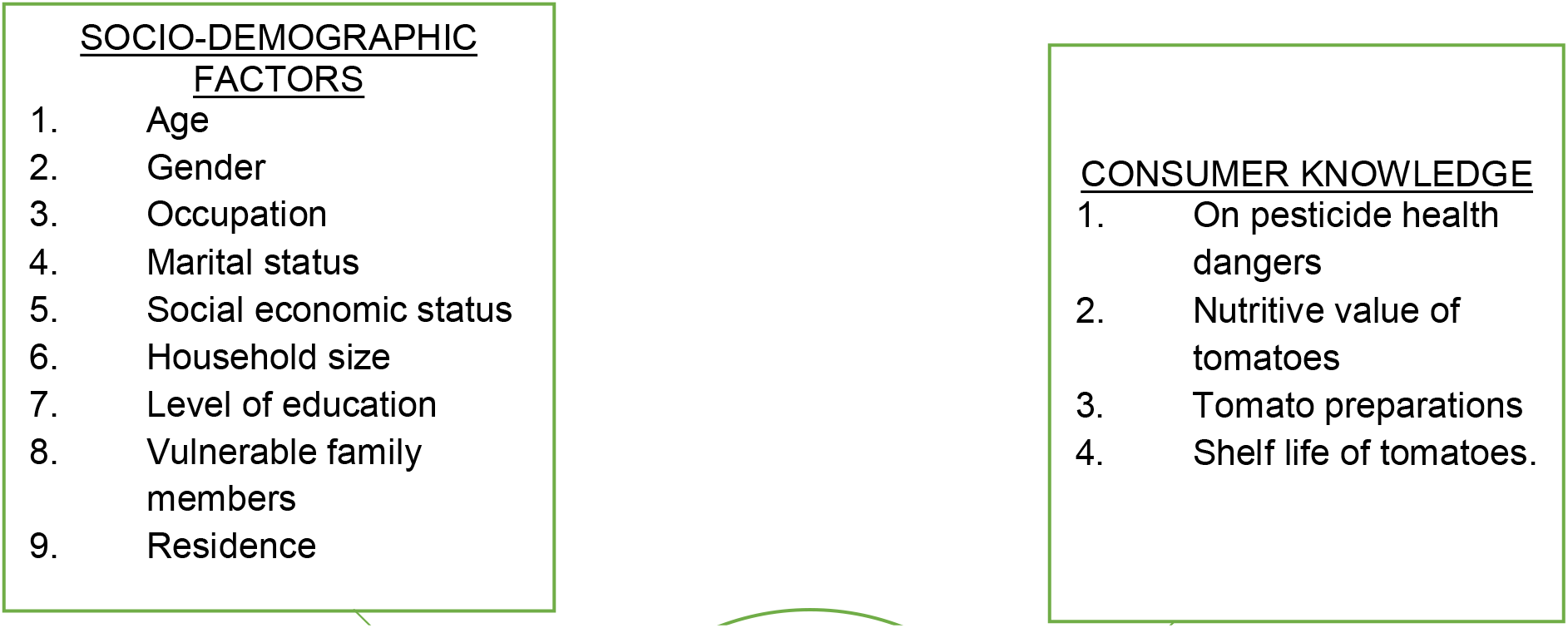

## Methods

### Study area and population

This study was conducted in 4 districts as interventional project areas, each selected from one of the four regions of Uganda namely, Northern (Nebbi), Western (Masindi), Eastern (Bugiri), and Central (Sembabule). From each district, three sub-counties were randomly selected and consumers were sampled systematically at the household level for interviews. The above districts were Pesticide Use, Health and Environment (PHE) Project intervention areas with a tropical climate where tomatoes, cabbages, passion fruits, oranges, mangoes, okra, green pepper, amaranths, and eggplant, among other crops, are commonly grown and intensively sprayed with pesticide. The commonly used method of pesticide application is a knapsack sprayer worn as a backpack and mechanically operated with a hand pump. The Uganda National Census 2014 estimates the average population for the above districts as follows; Nebbi (385,220), Masindi (94,622), Bugiri (426,000), and Sembabule (219,600). (29)

### Study design

A cross-sectional study design conducted in June 2019 employed mixed methods of both qualitative and quantitative data gathering. Working through existing branches of the District Farmers Association (DFA) in each of the four districts, three sub-counties were randomly selected and sampled, clustered into urban, peri-urban, and rural. In each of the four districts, an average of 117 participants at the household level, were randomly selected and interviewed making a total of 468 participants as follows; 117 participants from Nebbi District, 102 participants from Sembabule District, 119 participants from Bugiri District, and 130 participants from Masindi district. In addition, purposive sampling for Focus Group Discussions (FGDs) was conducted forming a total of 9 participants per focus group per district. Each FGD was composed of three tomato farmers, three tomato retail vendors, and three tomato wholesalers totaling 36 participants for the four districts.

## Materials

Pretested and standardized structured questionnaires adopted from a survey “*A monitor on consumer confidence in food safety*” developed (by Janneke de Jorge, 2008)(30) for monitoring consumer safety in a Canadian population were modified and used for data collection (details provided in the S5 File). All questionnaires were translated into the participant’s local language and translated back into English for quality assurance purposes. Focus Group Discussion Guides were used to collect the qualitative data and also administered in the local language by trained Research Assistants (RAs). (Details of the guides are provided in S6 File).

Consumer risk perception was assessed using a series of three questions in the order; 1) Are pesticide residues harmful to human health? 2) Are you aware that tomatoes sold on local markets contain pesticide residues? 3) Do you buy pesticide-stained tomatoes? Attitudes were measured on a three-Likert scale (responses ranging from agree, not sure and disagree). The questionnaire included three sections on optimism, pessimism, and trust. Under the optimism section, questions assessed the safety, confidence, and satisfaction of the pesticide residues on tomatoes. The section on pessimism assessed the worrisomeness, suspicion and discomfort caused by the pesticides residues on the tomatoes, while the section on trust assessed the consumer’s trust in whether the tomato vendors had the characteristics of trust such as the competence to control the safety of tomatoes, the knowledge to guarantee tomato safety, honesty about the safety of the tomatoes, sufficiently open about tomato safety and giving special attention to control the safety of tomatoes (Assessment results provided in S2 File).

### Data collection and analysis

Research Assistants (RAs) were trained on the objectives of the study in a one-day training per district. Questionnaires were pretested with the RAs, and supervision was done daily; every filled-in questionnaire was reviewed for accuracy and completeness to ensure data quality and ethical considerations were unbleached.

A total of 36 participants were involved in the FGDs, 9 per district, all their responses recorded using audio recorders and data gathered on tapes. The sample size for the FGDs was based on the level of saturation of the responses.

Quantitative collected data was gathered and entered into Microsoft Excel Version 2013 and exported into Stata version14 for cleaning and analysis. A total of 468 entries were achieved. Categorical variables like Risk Perception (measured as a binary outcome; high-risk or low-risk perception) and Attitude, age group, occupation, gender, and level of education were presented as frequencies with their respective percentages, while continuous variables such as age presented as means with their respective standard deviations (SD) and ranges.

Bivariable analysis for risk perception (measured as a binary outcome), was computed by gender, level of education, age categories, residence (rural, urban, and peri-urban), and the chi-square and the respective p-values reported under (95% CI, 5% Level of significance). Awareness about the pesticide residues was computed by level of education and by the practice of buying tomatoes and their Chi-square, p-values under (95% CI and 5% Level of significance) reported. Fisher exact test was used to determine the factors associated with consumer risk perception and the factors for buying pesticide-stained tomatoes under (95% CI, 5% Level of significance), details provided in the S3 File. Finally, simple logistic regression was used to compute the odds ratio of consumers who were aware of pesticide residues on the tomatoes vs those who were not aware of the pesticide residues on the tomatoes and their respective Odds Ratio with the p-values reported with 95% CI and 5% Level of significance as provided in the S4 File.

Qualitative data collected among the 36 participants in the four districts were transcribed and analyzed thematically based on the study objectives and conclusions based on participant responses. These conclusions in line with triangulation were later used to support a discussion with quantitative data findings.

### Ethical consideration

This study sought ethical approval from Makerere University School of Public Health, Higher Degree Research and Ethics Committee (MakSPH HDREC) with reference registration number 686. Informed consent was sought from all participants before the interviews; for anonymity, participants’ initials were used instead of their names on the questionnaire, and participants were free to withdraw from the study at any point when they felt like not continuing with the interviews.

## Results

### Demographic characteristics of consumers

The study registered all consumer responses, equally sampled by residence (rural, urban, and peri-urban). From the three sub-counties in each of the four districts of Uganda; (Northern region: Nebbi district, Eastern Region: Bugiri District, Central Region: Sembabule district, and Western region: Masindi district).

As indicated in Table 1 below, slightly more than half, 54.1% (253/468), of respondents were females, a majority, ≈51.0% (238/468) practiced farming as an occupation (*refer to S1* File for other demographic characteristics), a majority, 84.4% (395/468) had attained a lower level of education. The mean age of participants was 37.7 years (SD±13.1, ranging from 18-88), with a majority of 54.7% (256/468) belonging to the age group below the mean age. From the qualitative results, interviews involved categories of tomato farmers 33.3% (12/36), tomato retail vendors 33.3% (12/36), and tomato wholesalers 33.3% (12/36) sampled in equal proportions in the 4 districts. i.e., three persons, per category, per district.

**Table 1:** Demographic characteristics of consumers.

| Variable | Category | Frequency (%) |
| --- | --- | --- |
| <u>Gender</u><br>(n=468) | Male | 215 (45.9) |
|  | Female | 253 (54.2) |
| <u>Age group</u> | Below mean Age | 256 (54.7) |
|  | Above mean Age | 212 (45.3) |
|  | **Mean Age** | 37.7<br>(SD±13.1) |
| <u>Level of education</u> | No formal education | 39 (8.3) |
| (n=468) | Lower level | 395 (84.4) |
|  | upper level | 34 (7.3) |

### Consumer risk perception towards pesticide-stained tomatoes

Consumer risk perception was measured using a model with questions as provided in Fig 1.

**Fig 1.**
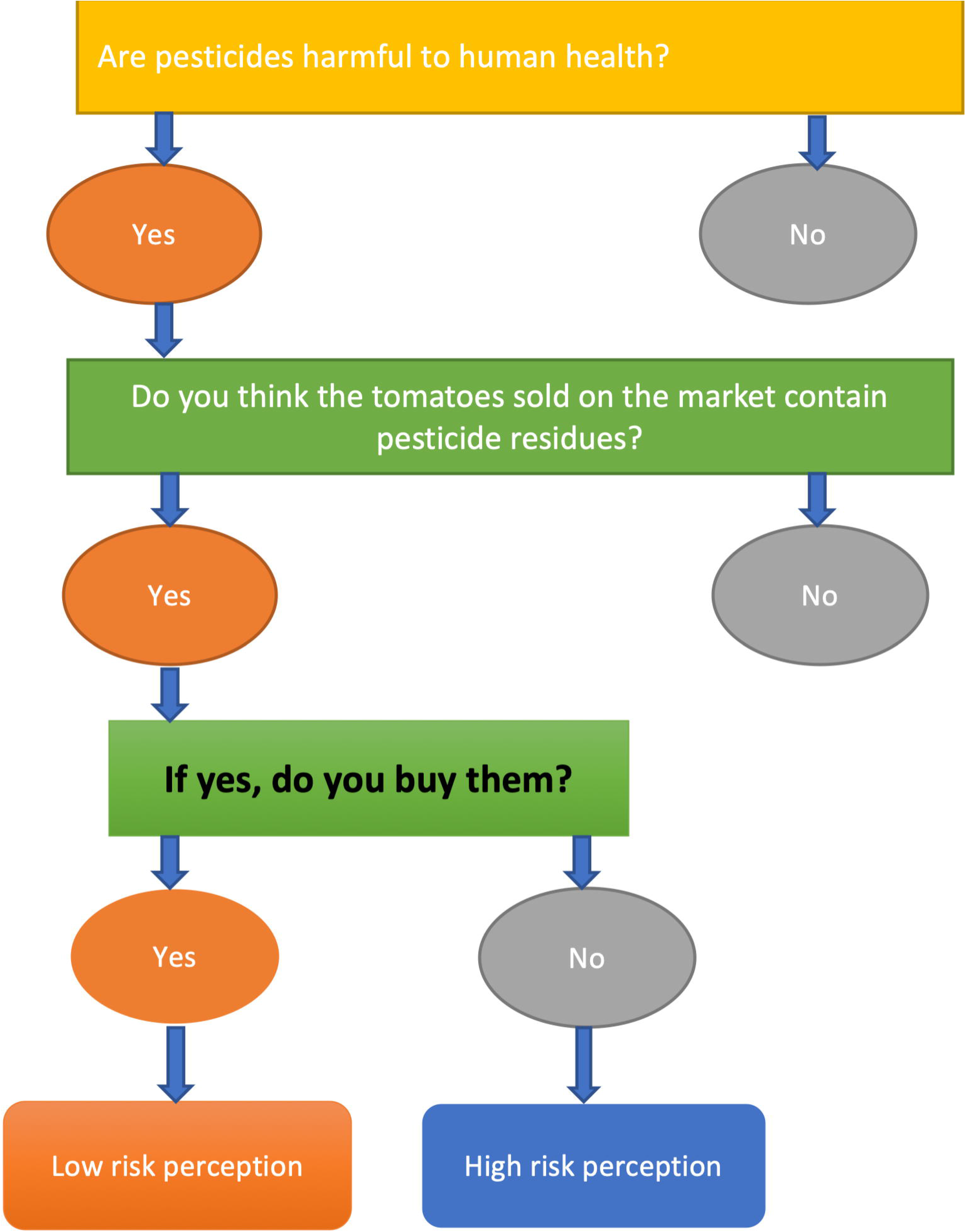
Model for determining consumer risk perception Consumer risk perception

Among consumers who were aware and knowledgeable about sold tomatoes containing pesticide residues, ≈95.0% (376/396) of them bought these pesticide-stained tomatoes (i.e., had a low-risk perception) compared to only 5.0% (20/396) who perceived tomatoes to be of high-risk to their health and withdrew from buying them (i.e., had a high-risk perception) as provided in Fig 2.

**Fig 2.**
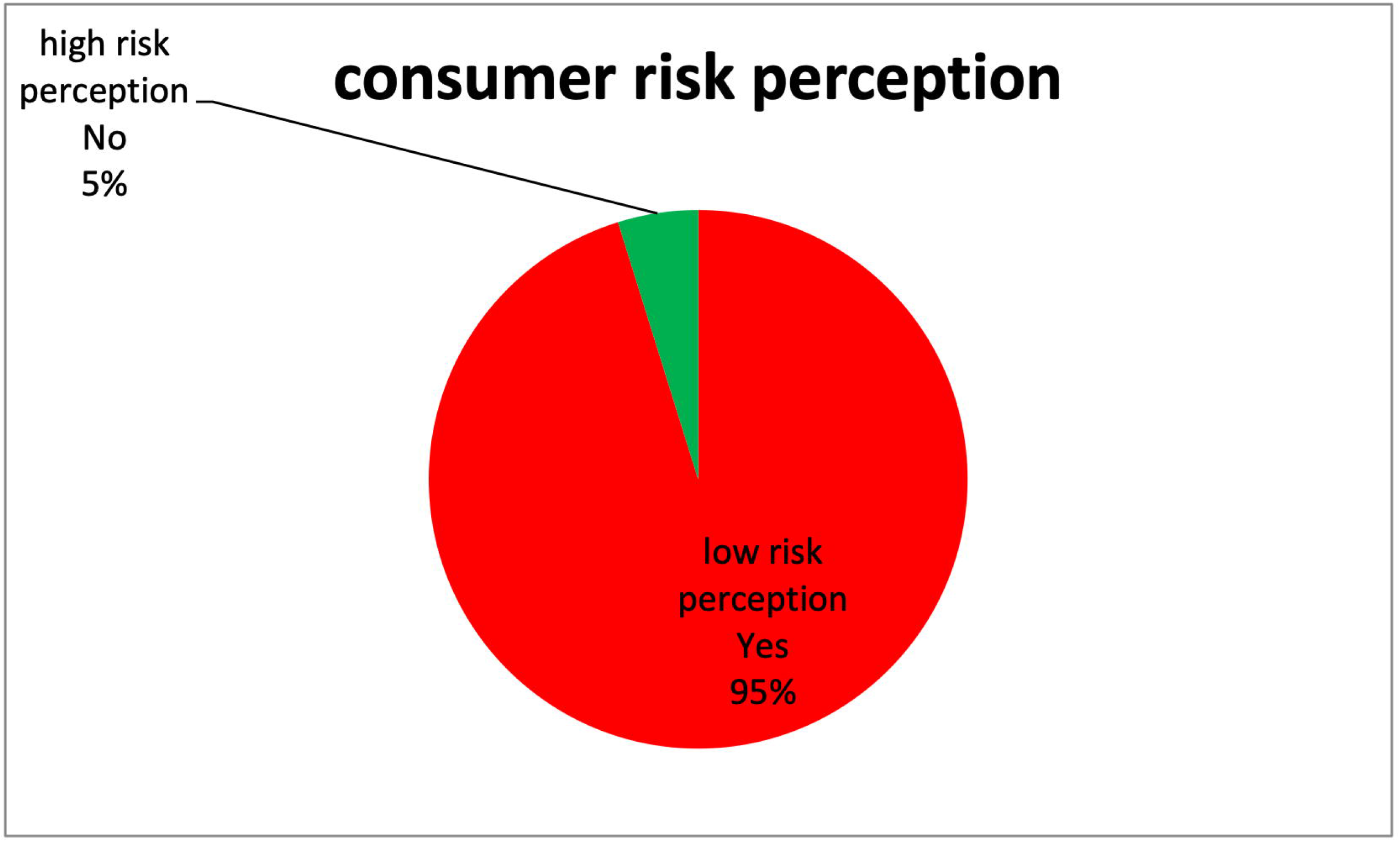
Level of consumer risk perception towards pesticide-stained tomatoes Reasons for buying pesticide-stained tomatoes.

**Fig 3.**
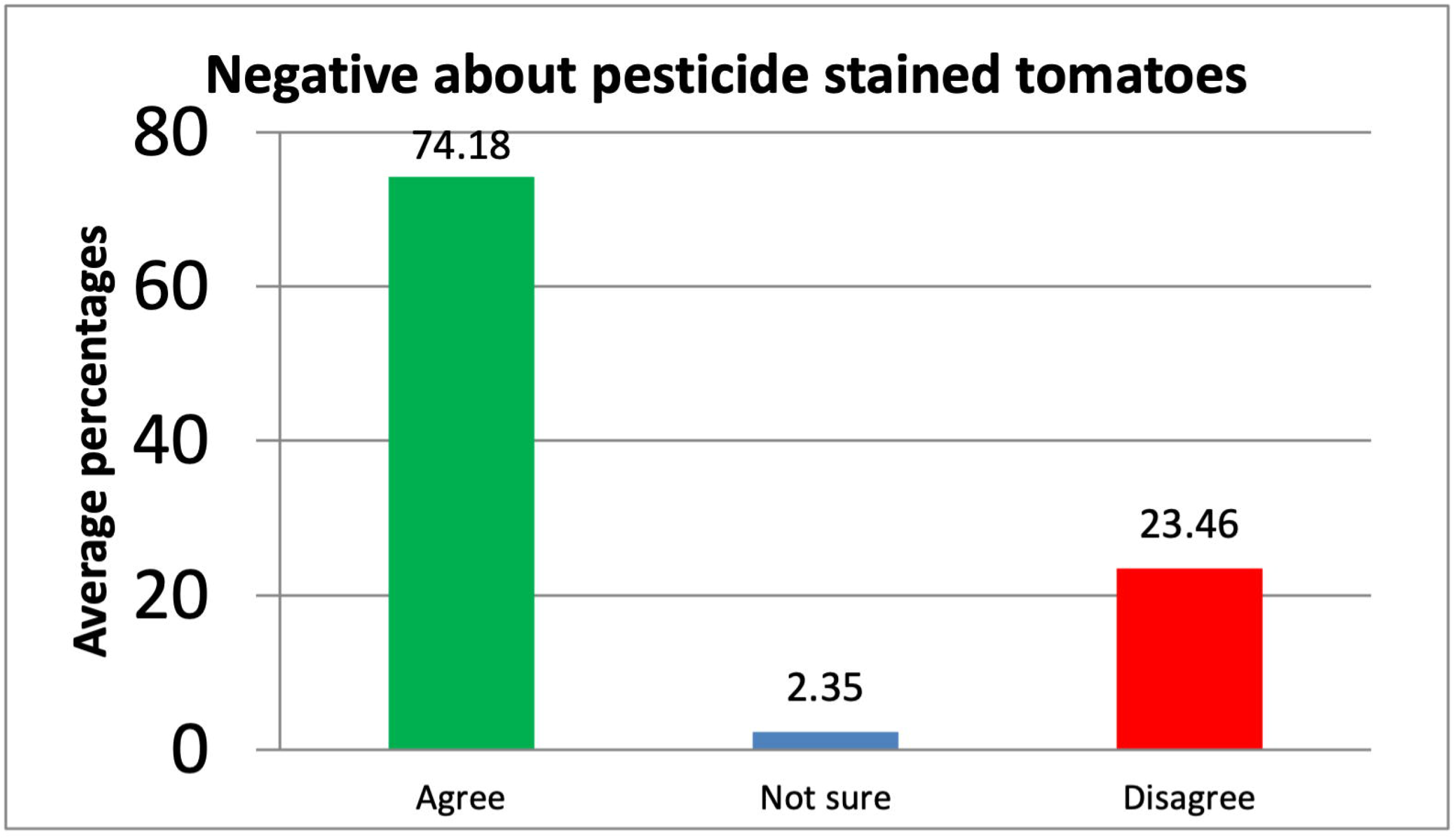
Average negative perceptions/pessimism of consumers on pesticide-stained tomatoes in percentage.

The main reasons for buying pesticide-stained tomatoes were reported that majority, 59.0% (230/390) of the consumers had no choice to buying pesticide-stained tomatoes, followed by 27.2% (106/390) who had to prepare these tomatoes at home to reduce the pesticide residues, 9.2% (36/390) who perceived no health risks of buying stained tomatoes and 4.6% (18/390) falling in the other categories which included the preference of tomato attributes like size, ripeness, price, among other attributes other than the pesticide residues.

### Consumer confidence in the safety of tomatoes sold in Ugandan Market

On a general scale, majority and more than half, 66.9% (313/468), of the consumers disagreed with the fact that tomatoes sold in the Ugandan markets are safe; consumers’ general confidence in the safety of tomatoes sold in Ugandan Markets outweighed their counterparts with nearly half, 49.6% (231/466), of the consumers being confident about the safety of tomatoes sold on the Ugandan markets. In comparison, only 14.4% (168/466) were not confident, and 14.4% (167/466) were not sure and 2 who never responded to the question.

### Factors associated with consumer risk perception and buying of stained tomatoes

From the Fisher-exact tests, consumer risk perception was not associated (p>0.05) with the demographics such as level of education P (0.975), Residence P (0.462), gender P (0.581), age-group P (0.680), and marital status P (0.581), as indicated in the S3 File. However, further simple logistic regression analysis revealed that consumer risk perception was significantly associated with consumer awareness about residues in the sold tomatoes; where the proportion of consumers who were aware of tomatoes containing pesticide residues was 42.8 times more of a high-risk perception compared to the proportion of those who were not aware of the tomatoes containing pesticide residues. (Refer to results in the S4 File).

In table 2, the consumer awareness about Pesticide residues was not associated with age-group and level of education but significantly associated with gender, where male consumers were 1.77 times more likely to be aware of the pesticide residues in the tomatoes compared to their counterparts; and consumers who had never obtained pesticide safety information being 61% less likely to be aware of the pesticide residues compared to consumers who had obtained information on pesticide safety, OR 0.39 (95% CI: 0.24-0.64)

**Table 2:** Logistic Regression for Consumer awareness about pesticide residues on tomatoes and some consumer demographics (crude Odds ratio)

|  | **Aware of pesticide residues** | **Odds Ratio** | **Std. Err.** | **P>\|z\|** | **[95% CI ]** |  |
| --- | --- | --- | --- | --- | --- | --- |
|  | **No (%) Yes (%)** |  |  |  |  |  |
| **Gender** |  |  |  |  |  |  |
| Female | 54(21.3)<br>200(78.7) | 1.0 |  |  |  |  |
| Male | 28(13.2) 184(86.8) | 1.77 | .451 | **0.024** | 1.078 | 2.921 |
| **Age category** |  |  |  |  |  |  |
| Below mean age | 46(18.0) 210(82.0) | 1.0 |  |  |  |  |
| above mean age | 36(17.1) 174(82.9) | 1.06 | .259 | 0.816 | .655 | 1.711 |
| **Education category** |  |  |  |  |  |  |
| None | 10(25.6) 29(74.4) | 1.0 |  |  |  |  |
| Lower-level Education | 69(17.6)<br>324(82.4) | 1.62 | .631 | 0.217 | .754 | 3.477 |
| Upper-level Education | 3(8.8) 31(91.2) | 3.56 | 2.52 | 0.072 | .891 | 14.249 |
| **Ever obtained information about pesticide safety** |  |  |  |  |  |  |
| Yes | 34(12.1)<br>247(87.9) | 1.0 |  |  |  |  |
| No | 48(25.9)<br>137(74.1) | 0.39 | .098 | **0.000** | .244 | .639 |

Consumer awareness about the pesticide-stained tomatoes was not associated with consumer level of education P (>0.05) but significantly associated with consumer risk perception P(<0.05) and the practice of buying stained tomatoes P (<0.05) as indicated in Table 3 below.

**Table 3.** shows the fisher-exact tests for awareness about pesticide residues in tomatoes versus the consumer level of education and practice of buying stained tomatoes.

| **Buy stained tomatoes** | **Aware of pesticide residues in the tomatoes Freq (%)** |  |  | **Fishers-exact p-values** |
| --- | --- | --- | --- | --- |
|  | **Yes** | **No** | **Not sure** | **0.000** |
| No | 14/20(70.0) | 6/20(30.0) | 0 /20(0.0) |  |
| Yes | 370/376(98.4) | 4/376(1.1) | 2/376(0.5) |  |
| **Consumer risk perception** |  |  |  | **0.000** |
| High-risk perception | 13/19(68.4) | 6/19(31.6) | 0/19(0.0) |  |
| Low-risk perception | 371/377(98.4) | 4/377(1.1) | 2/377(0.5) |  |
| **Education level** |  |  |  | 0.095 |
| None | 29/39(74.4) | 7/39(17.9) | 3/39(7.7) |  |
| primary | 216/272(79.4) | 35/272(12.9) | 21/272(7.7) |  |
| secondary | 108/121(89.3) | 7/121(5.8) | 6/121(5.0) |  |
| tertiary | 31/34(91.2) | 1/34(2.9) | 2/34(5.9) |  |

### General consumer attitudes towards pesticide-stained tomatoes Pessimism towards pesticide-stained tomatoes

Based on the percentages of pessimism, measured on a 3-Likert scale, a majority, 74.3% (347/467) of the consumers were pessimistic about the stains on the tomatoes compared to 2.4%(11/467) who were not sure and 23.5% (109.6/467) who felt optimistic about the stains on the tomatoes in terms of worrisome, discomfort, and suspicion caused by the residues. (Details provided in S2 File and Fig3 ).

### Optimism towards pesticide-stained tomatoes

Based on the percentages of optimism, measured on a 3-Likert scale, Consumers’ positive attitude towards pesticide-stained tomatoes was low. On average, only 33.2% (155.3/468) of the consumers agreed with the statement that tomatoes sold on the Ugandan market are safe, compared to a majority of 61.1% (285.6/468) who disagreed with the statement while 5.7% (26.7/468) were not sure as provided in Fig 4. and indicated S2 File.

**Fig 4.**
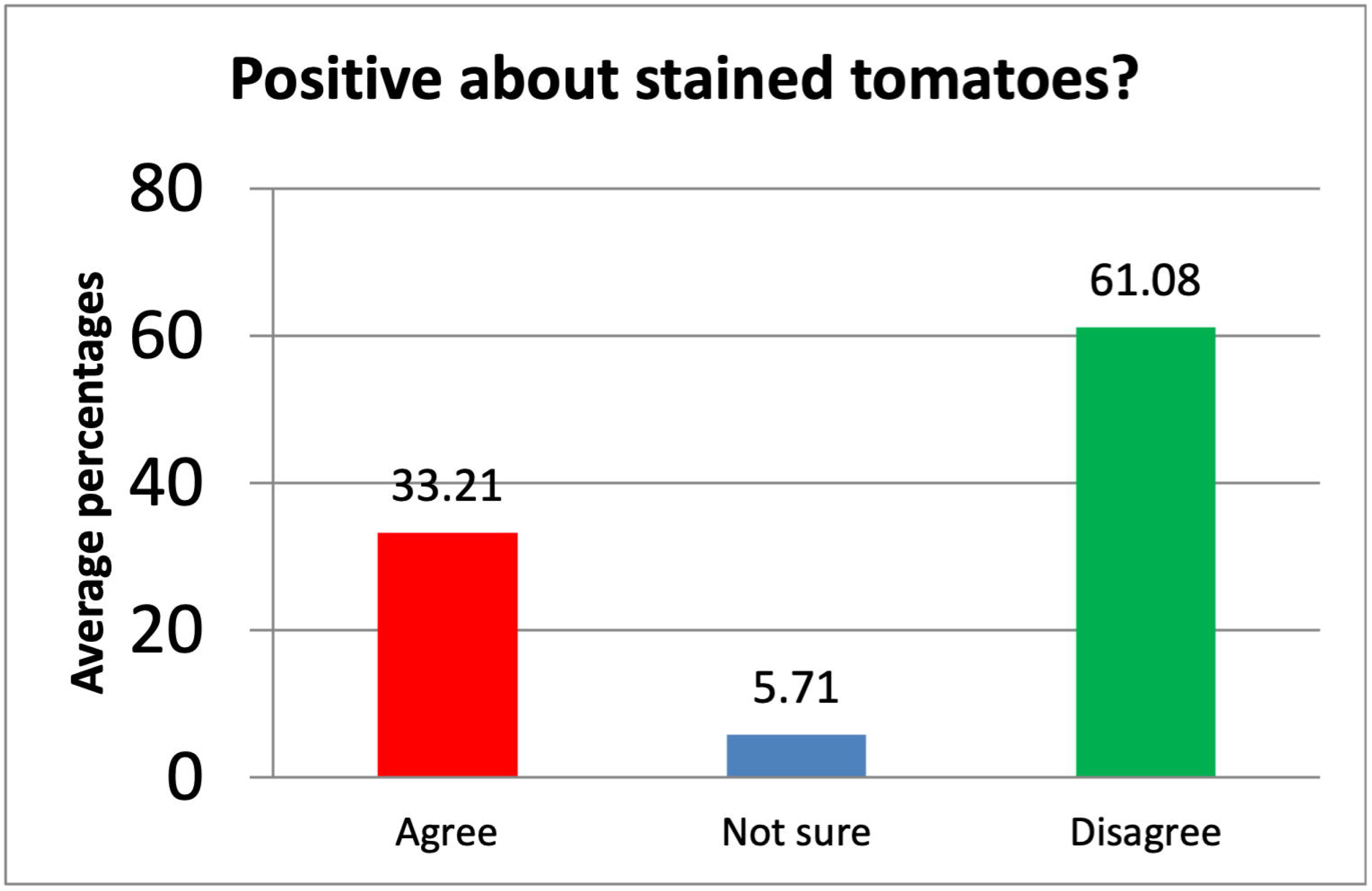
The optimism of consumers on pesticide-stained tomatoes.

### Consumer-level of trust in pesticide-stained tomatoes

In terms of trust, based on the 3-scale, 77.7 % (362/466) of consumers lacked trust and disagreed that tomato vendors have the characteristics of trust such as the competence to control the safety of tomatoes, the knowledge to guarantee tomato safety, honesty about the safety of the tomatoes, being sufficiently open about tomato safety and giving special attention to control the safety of tomatoes compared to 8.1% (37.6/466) who were not sure and 14.1% (65.4/466) who trusted tomato vendors (details provided in the S2 File).

### Qualitative findings

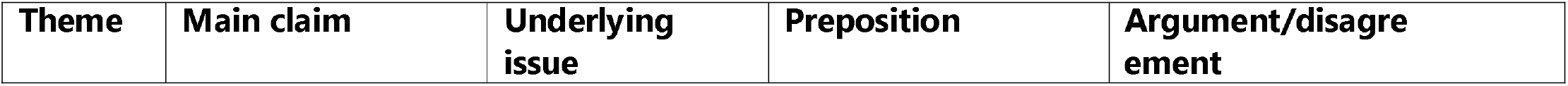
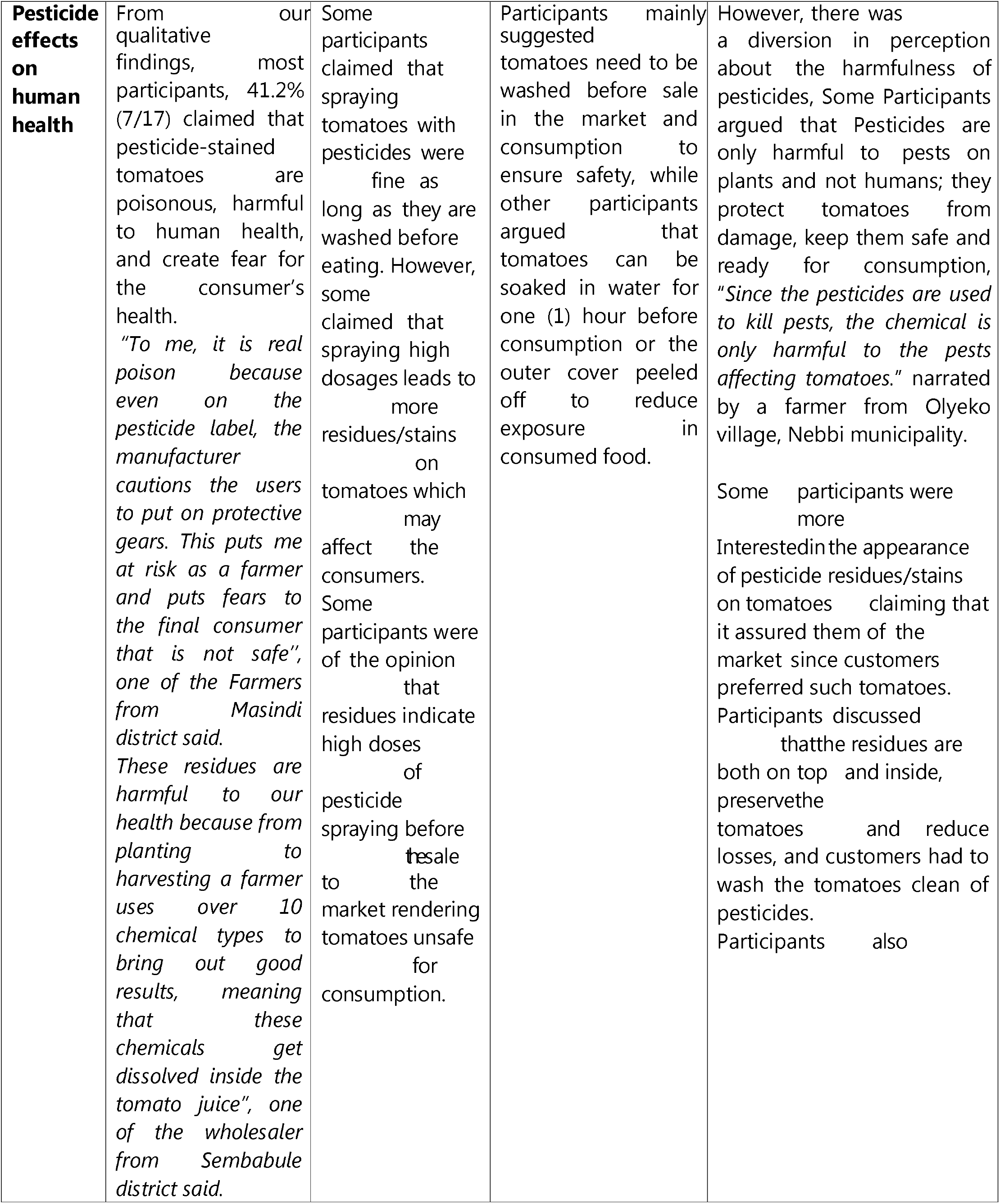
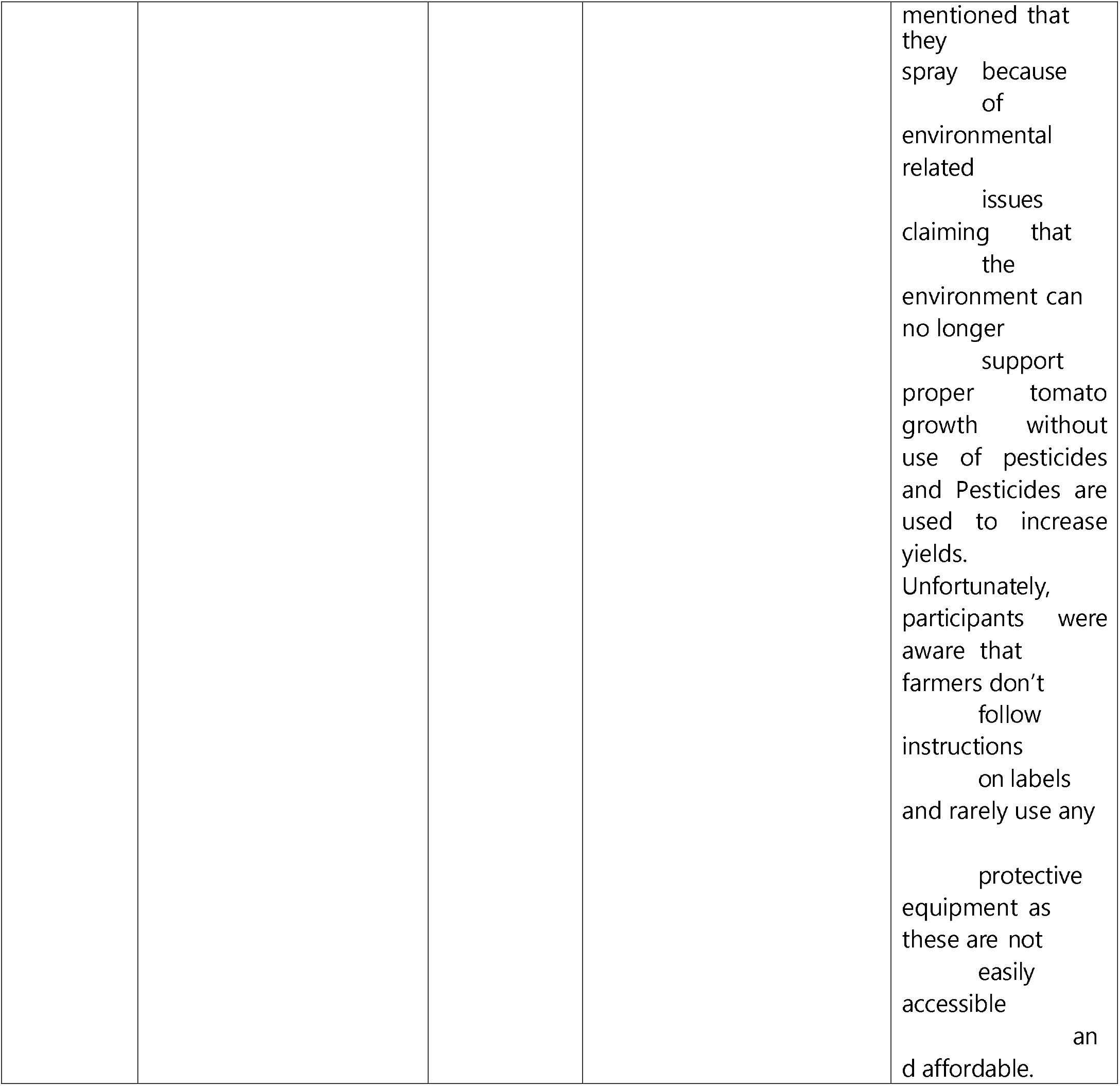
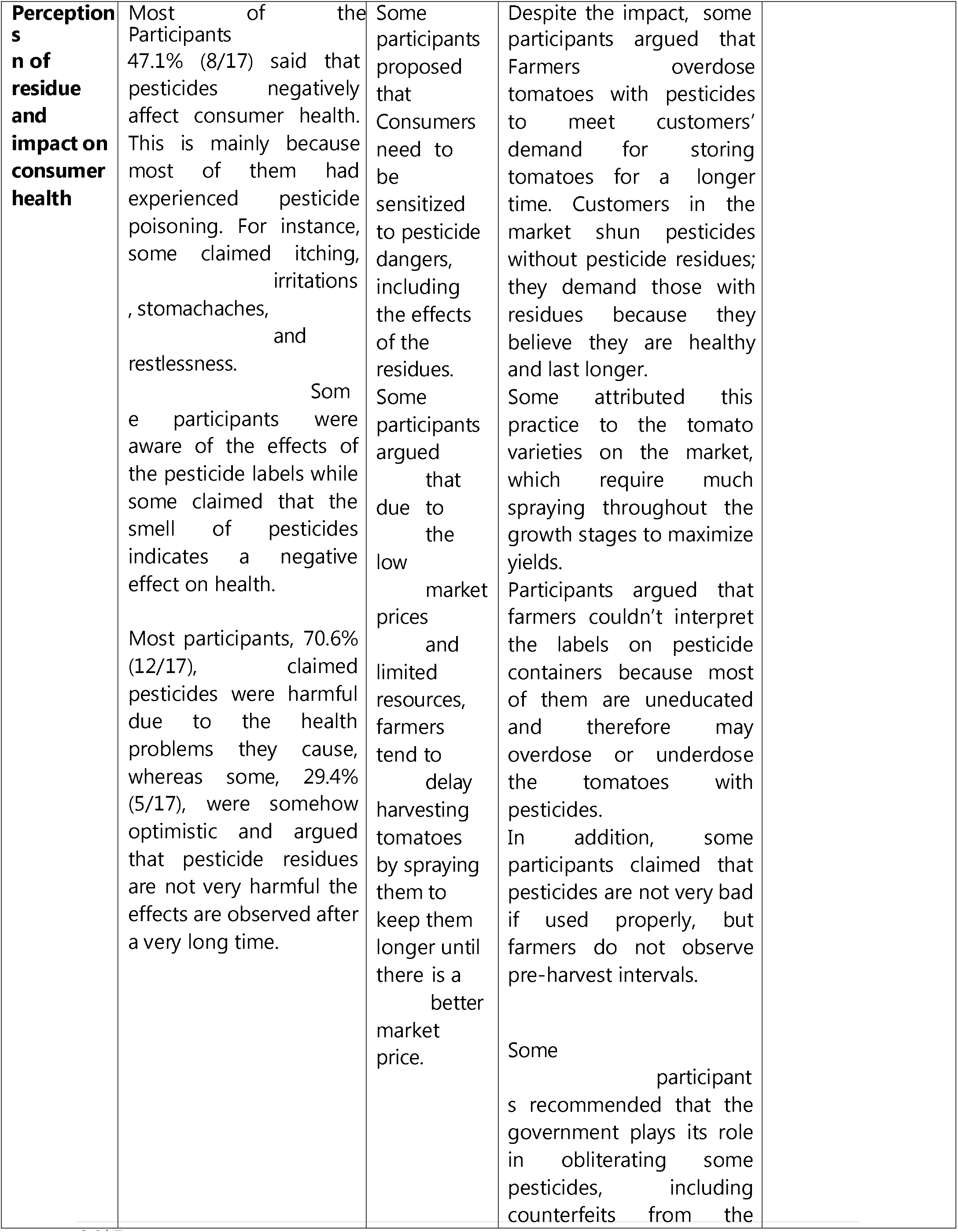
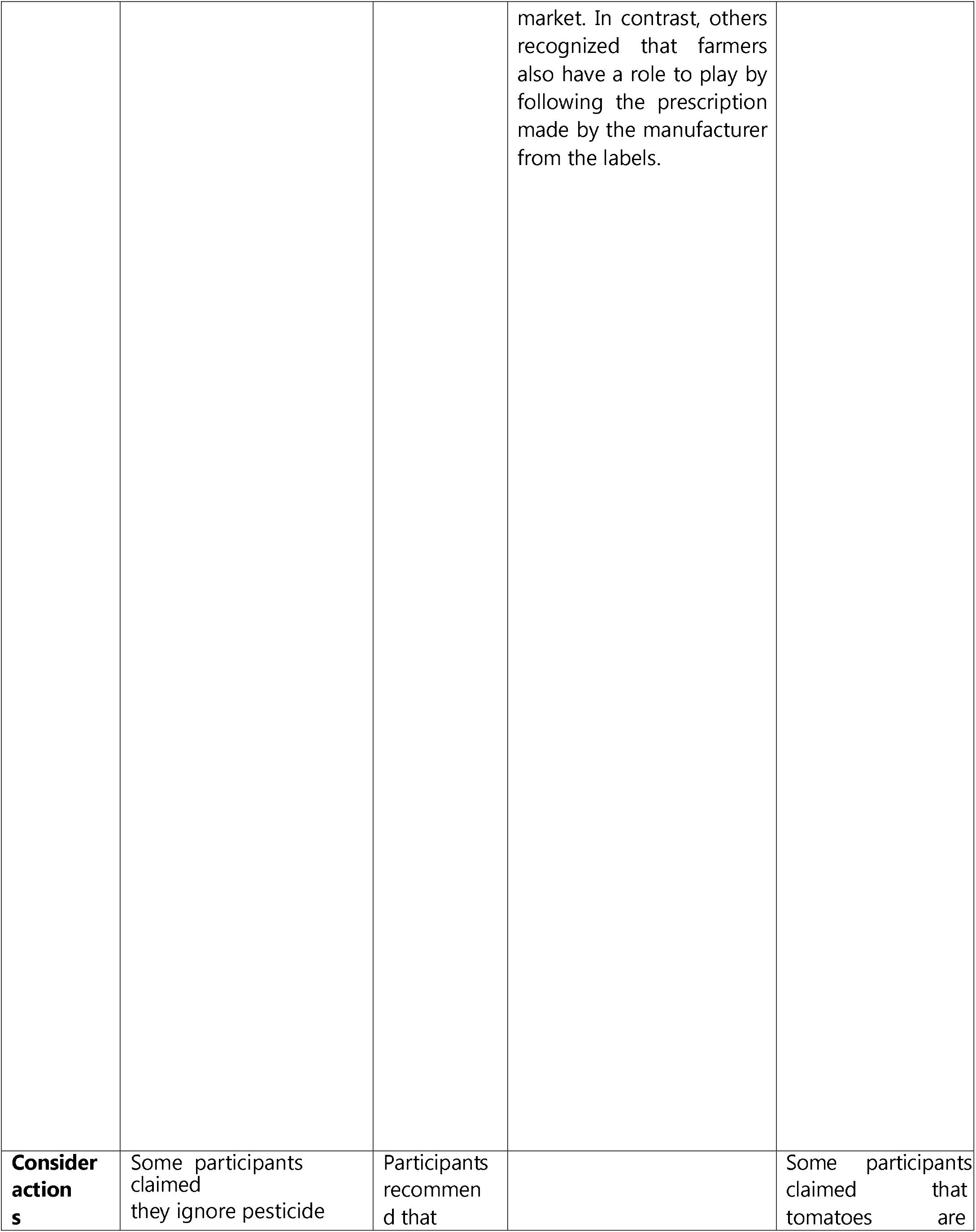
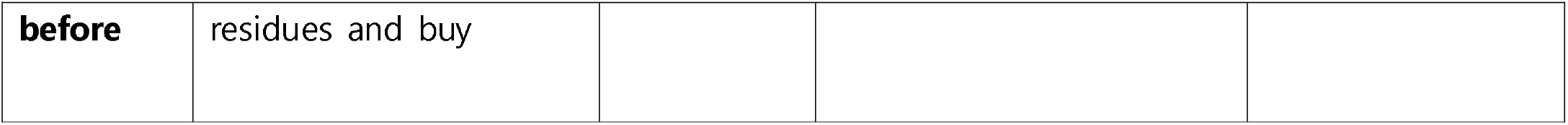
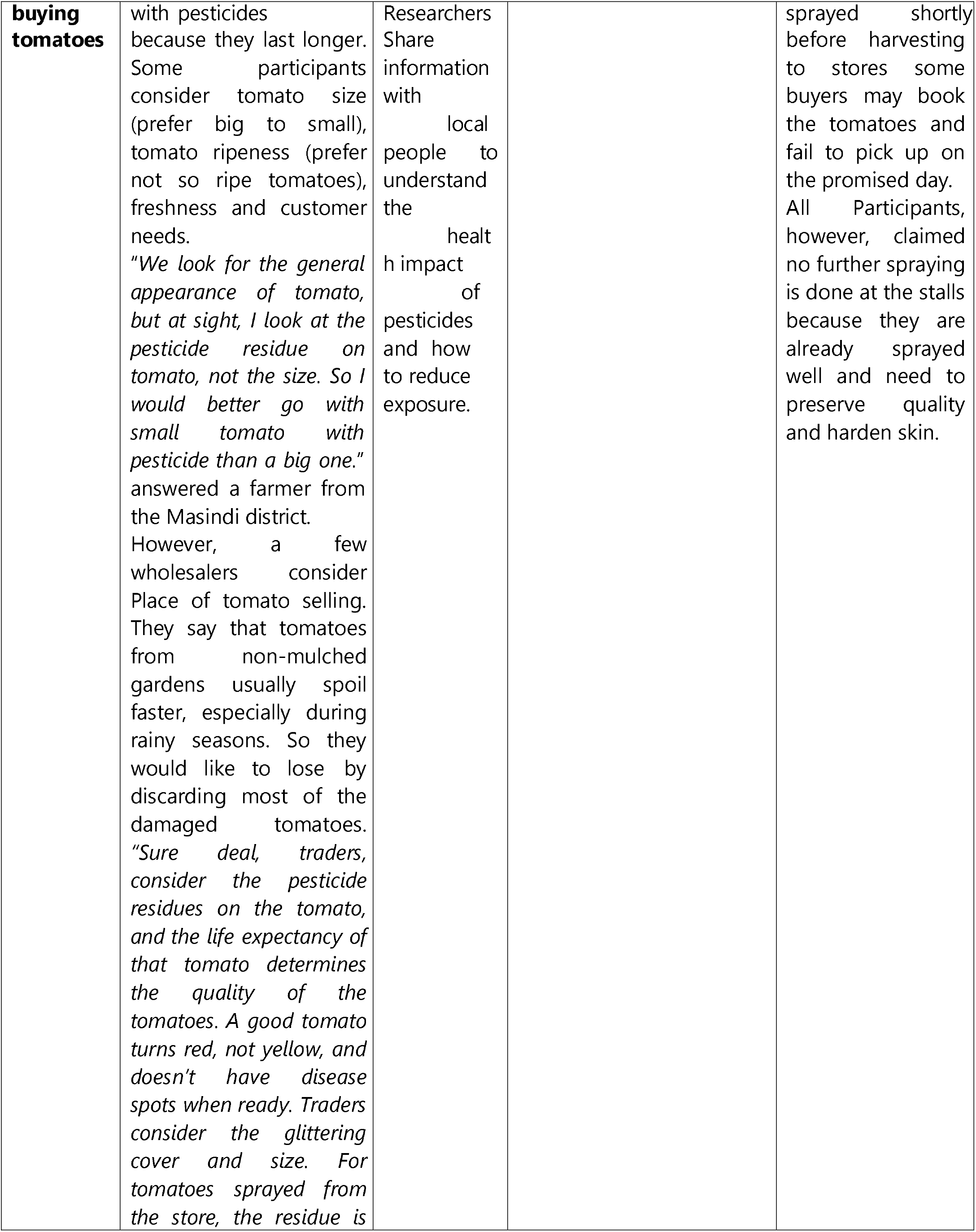
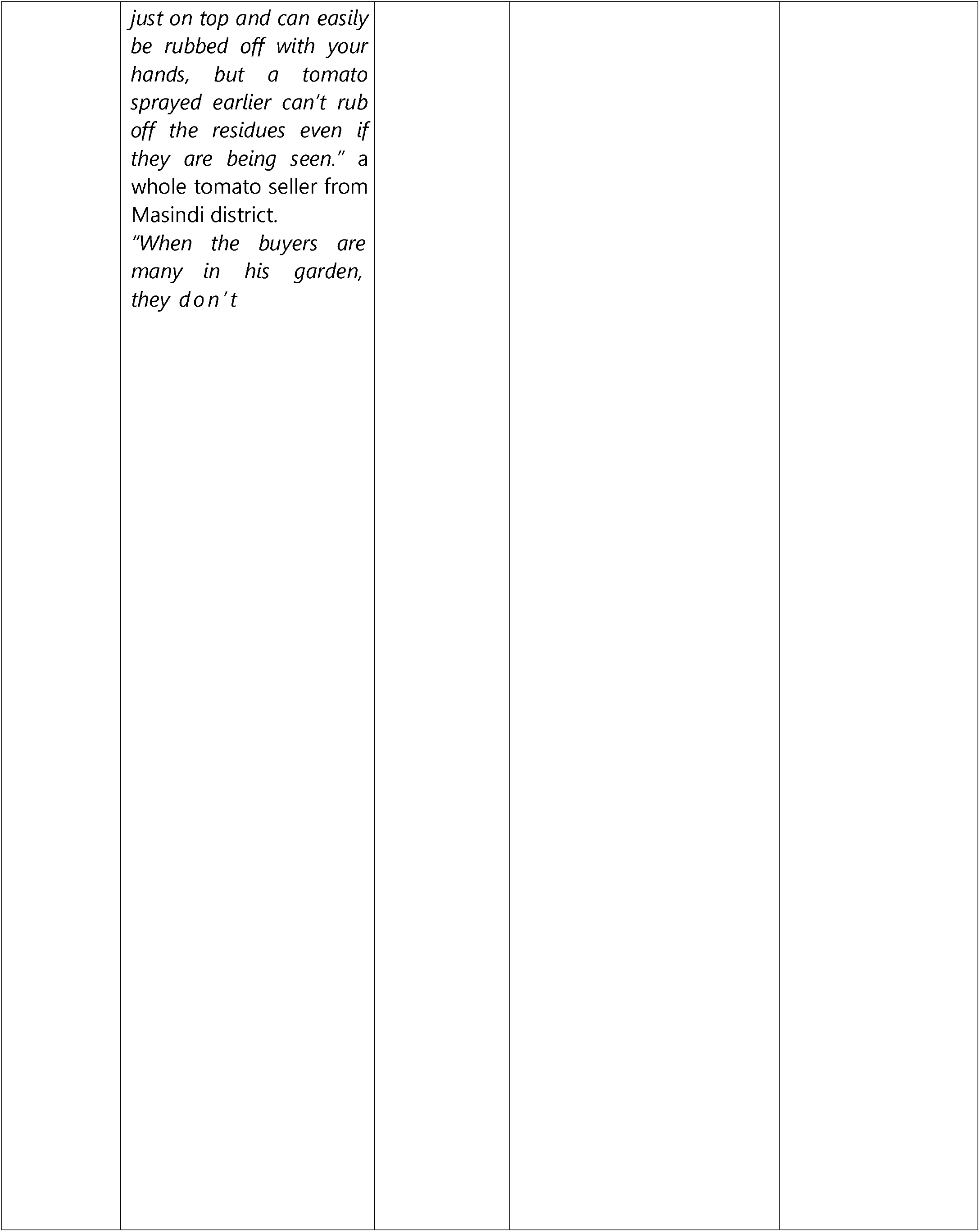
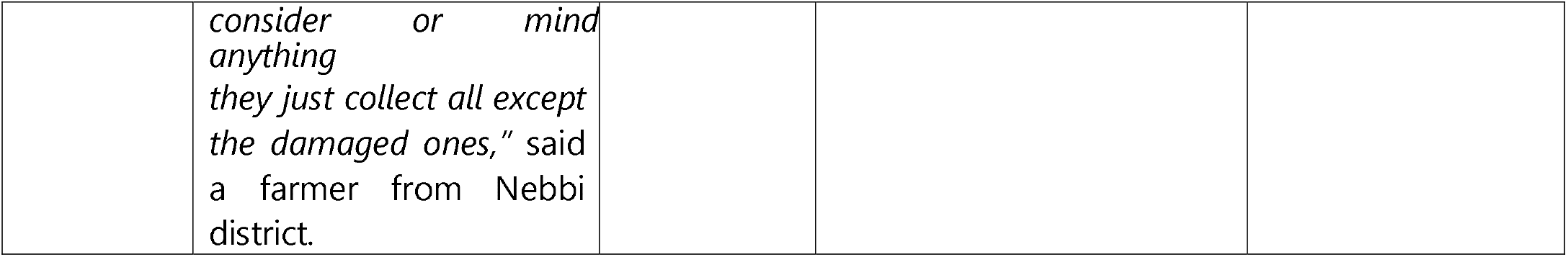

## Discussion

Very little information is available about the pesticide residue in tomatoes in Uganda and, consequently the risk of exposure to this. Therefore, the focus of this study was to determine the consumer risk perception towards pesticide-stained tomatoes and the attitudes (pessimism, optimism and trust) toward pesticide-stained tomatoes. It employed a cross-sectional study design with a total of 468 consumers as respondents equally systematically sampled by residence and interviewed from each of the four districts (Northern region: Nebbi district, Eastern Region: Bugiri District, Central Region: Sembabule district, and Western region: Masindi district), thus a good representation of Ugandan consumers.

Characteristics of the respondents show that slightly more than half of the respondents were females as expected since data collection was done at home level and majority of the females stay home to take care (31) of home chores and are more involved in food buying and preparations, similar to findings by (32, 33) where 58% of women were involved in the purchase of meat and (34) where 73% of the female formed part of the study participants, more than three quarters were married women as interviews were conducted at homesteads and consent sought from adults making it more likely for the married women to be interviewed. This calls for public health and risk communication interventions to focus and prioritize females where communicating the risks of pesticides exposure in food since they are more engaged in food preparations at home levels. Half of the respondents were farmers, given that Uganda has nearly three-quarters of its population engaged in farming, and this study was carried out in a rural setting involving vendors, buyers, and tomato growers. A majority had completed lower level of Education, and a majority belonged to the age group <30 years as the Ugandan population is composed of 75% youths as the majority, 80% of these residing in rural areas (35). From these characteristics, we can report that the consumers interviewed in this study were a good representation of tomato consumers in Uganda. Respondents of this study represent adult female consumers experienced with tomato farming and with adequate education attainment (majority with Ordinary level of education) to express risk perception towards pesticide-stained tomatoes.

Consumer risk perception ranked low, with a majority of consumers (95%) buying tomatoes that are well known to be stained with pesticide residues, a majority giving reasons that they have no alternative. Results from this study highly deviate from similar studies conducted in developed countries, such as Californian consumers, where 80% were safety cautious and checked the food items to see if they were opened or damaged (36), in Georgia, 89% considered testing of pesticide residues in food to be very important or somewhat necessary (31), and in Boston, consumers had a high-risk perception of conventionally grown produce compared to public health hazards (37), most of these were mainly triggered by the health effects of the contaminants such as pesticide residues and biological contaminants like disease causing germs in the sold produce. More so, another study by (34) conducted in Athens Greece, shows how consumer willingness to pay for organically grown food was strongly linked to quality and security. Quality entails the aspects of food being free from any contaminations introduced in the process of its growth and the security aspect entailing a low risk to harm.

From the Turkish perspective, consumers’ willingness to pay for reduced pesticide residues in tomatoes was mainly determined by their risk perception about the residues, which is explained by the label on the purchased apples (38). In this case, due to the different situations, in Uganda, unfortunately, tomatoes are not labeled with the residual contents and benefits of low pesticide residual levels. This could be the reason why most of the consumers in this study bought tomatoes. The vegetables on the market lack such information which could have triggered consumers to make their choices. This is a gap to be closed by authorities in charge of food safety and consumer protection to ensure that food is labeled in terms of chemical residues but also state if it is organically grown to protect the public from pesticide residual exposures. Uganda is lacking a stand-alone and comprehensive food safety policy that puts public health at stake for pesticide exposure. The existing Food and Drug Act is a bit outdated and merely captures specific food contaminants such as pesticide residues.

Results from this study are a piece of vivid evidence to be used as part of the advocacy statements in finding ways of establishing a stand-alone National Food Safety Policy and formulation of interventions for risk communication regarding food risks like pesticide residues.

A study on consumers’ willingness to pay for pesticide-free vegetables indicated how consumer awareness about the residues in vegetables and the residual effects on human health greatly influenced their willingness to pay for these vegetables. Consumers in this study were willing to pay 50% more for pesticide-free vegetables (39). In the Ugandan context, the consumer’s low-risk perception of pesticide-stained tomatoes indicates a risk of increasing pesticide residues exposure if there lacks an alternative. As reported in other findings (19), our qualitative results from vendors and tomato farmers from the FGDs, indicate that the highly stained tomatoes are due to poor hybrid tomato seeds that need frequent spraying and vendors’ demand to spray these tomatoes before selling. Nevertheless, a low level of literacy to understand the pesticide label information and wrong perception that Mancozeb pesticide can harden the outer skin and increase tomato shelf life. From this misconception, tomato vendors only buy stained tomatoes presuming that these will stay for long on the shelf, are healthy & free from microbial contaminants.

Unlike other studies (31) from Georgia where consumers prioritize microbial contamination followed by pesticide residues as the first consideration before choosing to buy vegetables, tomato consumers in Uganda partly have a feeling that pesticide-stained tomatoes are free from pathogens and healthy, thus ignoring the pesticide stains on them. As reported, some think that pesticides are selective and only meant to kill plant pests and cure plant diseases. However, our logistic regression analysis indicates how consumer awareness about pesticide residues increases the chances of a high-risk perception, protecting consumers from residue exposure, although this is just for a few individuals, this is backed up by the risk perception consumer behavior theory under perceived severity of the risk associated with the choices under the conventionally grown foods. More interventions on tomato preservation techniques need to be disseminated by district agricultural extension departments targeting more tomato sellers who demand pesticide-stained tomatoes.

From our focus group discussion; all farmers claimed that vendors would only buy tomatoes with pesticide stains as these are thought to be healthy and would stay for long on the shelf before being sold to the consumers. On the other hand, vendors attest that stained tomatoes are healthy, look good, take long to go stale and have a high resale value on the market.

From these findings, the Ministry of Agriculture Animal Industry and Fisheries needs to sensitize farmers and improve coordination and regulations on the sale and use of agro-inputs. Agro-input dealers, the immediate information providers to the farmers, need to be trained in Pesticide safe-use training. A recent unpublished survey done by UNACOH in 2020 reports that only 6% of the agro-input dealers in 12 districts had obtained the safe-use training which is required to be undertaken by all agro-input distributors and it is a prerequisite by law before an agro-input shop business is opened (40).

Tomato residues are given less attention by consumers probably because consumers lack knowledge of the dangers that the residues may impact on their health which in most cases takes time. From a model by Huang Chung (41) estimating the relationship between consumer perceptions, attitudes, and behavioral intentions (refer to the model in S7 File), choices (behavioral intentions) on buying food are influenced by perceptions and attitudes, which influence each other in addition to knowledge (information) from personal experience, evaluative criteria, and social demographics. All are based on consumer awareness about the pesticide’s potential ill effects and the social economic status of the consumer. Consumers of higher social-economic status are most likely to have a high educational level and consequently easily access all the necessary information about the effects of pesticide residues. These will tend to have a high-risk perception about the pesticide-stained tomatoes and are not likely to buy those stained with pesticides. However, this was not so with our findings. Consumer risk perception was not directly associated with the level of education and social economic status, but associated with awareness about pesticide residues. Consumers who were aware of pesticide residues were 42.8 times more likely to be of high-risk perception than those who were not aware. From our results, the low proportions (5%) of high-risk perception consumers may be attributed mainly to our sample containing low percentages of highly educated consumers (upper-level education, table 1). This study is in line with (34) discovery that consumer willingness to pay for organic produce was not associated with social demographic characteristics. However, similarly, consumer willingness to pay dwelt much on the quality and safety of the food. Where consumers look for quality food free from contaminants and this is affected by their knowledge or awareness about the contaminants.

On the other hand, pesticide residue knowledge among the general public in Uganda is a new topic, and studies conducted along these lines are few, most of the time not intended to create awareness among the public on the potential ill effects of residues in food. From other related findings in this study, consumers have no access to sources of information on pesticide residues in food, with most of the information on pesticide residues acquired through radio and television media, followed by health professionals, depicting a significant gap in information accessibility and availability. Based on the results of the misconception that pesticide residues increase tomatoes’ shelf life, new studies on the origin of this misconception need to be conducted to challenge the practices of overdosing, which pose risks to public health.

Although quality assurance measures were put in place and followed by research assistants to ensure that all reported and recorded data from respondents was the truth and nothing but the truth. Like all studies, this study faced limitations which included recall bias; for questions that required respondents to report through recalling food choices made the previous month or weeks, it’s expected that not all can report accurate information (report bias) but also questions relating to whether they consume pesticides contaminated food seems a sensitive matter, especially to respondents who are aware of dangers related to consumption of food contaminated with pesticide. Report bias is another limitation of this research.

## Conclusion

Although consumers in Uganda had a negative attitude towards pesticide residues on the tomatoes, their risk perception towards these pesticide-stained tomatoes ranked low, with a majority of consumers buying pesticide-stained tomatoes regardless of their level of education, age, and gender. This was all linked to a market lack of alternative organic tomatoes. However, awareness about tomatoes containing pesticide residues was associated with containing pesticide residues was 42.8 times more of a high-risk perception than consumers who were not aware.

There is a need by the government, through its line of Ministry of Agriculture Animal Industry and Fisheries (MAAIF), health information and risk communication entities, and Civil Society Organizations among other partners to sensitize the Ugandan population on the effects of pesticide residues. There is also a need to sensitize farmers on the right pesticide dosage, and compliance to the pre-harvest intervals as well as train agro-input dealers on the same relevant pesticide safety and handling modules since it is from these, that the farmers buy pesticides. MAAIF should also hasten the establishment of the National Pesticide Residue Monitoring Program to protect Public Health from this chronic exposure to pesticide residues in agricultural produce.

## Acknowledgments

The author would like to acknowledge the support from Diálogos participants and all efforts rendered by the Pesticides Use, Health and Environment (PHE) Project team, and the respective District Framers’ Association with which the study was conducted. DFAs supported highly in questionnaire translation into the respective local languages and in data collection.

## Supportive information

Data-set used to yield the above results is available upon request from the author. However, separate supportive information has been attached, as highlighted below.

S1 File. Other Demographic characteristics of consumers. PDF

S2 File. Consumer attitudes towards pesticide-stained tomatoes. PDF

S3 File. Fisher-exact tests for the factors associated with consumer risk perception towards pesticide-stained tomatoes. PDF

S4 File. Simple logistic regression of consumer risk perception vs. awareness about pesticide residues. PDF

S5 File. Questionnaire: Consumers’ risk perception towards pesticide-stained tomatoes in Uganda. PDF

S6 File. Focus Group Discussion Guide for Consumer risk perception towards Pesticide-stained tomatoes in Uganda. PDF

S7 File. A model for estimating the relationship between consumer perceptions, attitudes, and behavioral intentions was adopted from Huang Chung. PDF

## Notes

### Competing Interest Statement

The authors have declared no competing interest.

### Summary of Updates

the whole manuscript has been uniformly revised including a section that forms a basis for the methodology and the theory model for consumer perception added.

